# A taste for numbers: *Caenorhabditis elegans* foraging follows a low-dimensional rule of thumb

**DOI:** 10.1101/2022.06.21.496406

**Authors:** Gabriel Madirolas, Alid Al-Asmar, Lydia Gaouar, Leslie Marie-Louise, Andrea Garza-Enriquez, Mikail Khona, Christoph Ratzke, Jeff Gore, Alfonso Pérez-Escudero

**Affiliations:** Research Centre on Animal Cognition (CRCA), Centre for Integrative Biology (CBI), Toulouse University, CNRS, UPS, Toulouse 31062, France; Physics of Living Systems Group, Department of Physics, Massachusetts Institute of Technology, United States; Interfaculty Institute for Microbiology and Infection Medicine Tübingen (IMIT), Cluster of Excellence EXC 2124 “Controlling Microbes to Fight Infections” (CMFI), University of Tübingen, Calwerstrasse 7/1, 72076 Tübingen

## Abstract

Rules of thumb are behavioral algorithms that approximate optimal behavior while lowering cognitive and sensory costs. One way to reduce these costs is by reducing dimensionality: While the theoretically optimal behavior may depend on many environmental variables, a rule of thumb may use a low-dimensional combination of variables that performs reasonably well. Experimental proof of a dimensionality reduction requires an exhaustive mapping of all relevant combinations of several environmental parameters, which we performed for *Caenorhabditis elegans* foraging by covering all combinations of food density (across 4 orders of magnitude) and food type (across 12 bacterial strains). We found a one-dimensional rule: Worms respond to food density measured as number of bacteria per unit surface, disregarding other factors such as biomass content or bacterial strain. We also measured fitness experimentally, determining that the rule is near-optimal and therefore constitutes a rule of thumb that leverages the most informative environmental variable.

## Introduction

Sophisticated and highly optimal outcomes of animal behavior often emerge from simple rules, called rules of thumb.^1–7^ For example, some ants seem to be able to measure the area of a potential nest, yet they are in fact implementing a simpler algorithm: They move randomly inside the potential nest, counting the number of intersections with their own path (which is marked with pheromones). This number of intersections correlates inversely with the area, so this simple rule of thumb provides a near-optimal response while being easy to implement.^2^ Identifying these rules of thumb is key to link the neural and mechanistic implementation of animal behavior to the selective pressures that shape it.^8^

While most of the rules of thumb studied so far focus on simplified mechanisms to acquire and process information, decisions can also be simplified by lowering the dimensionality of the input variables. For example, optimal food choice may require considering simultaneously many variables such as the density of each food source, its composition in terms of many different nutrients, the spatial distribution of different food sources, etc. Processing all these variables separately is costly, so a rule of thumb may disregard the less informative variables and combine the rest into one or a few quantities that determine the decision. While numerous studies identify variables that dominate behavior^9^, demonstrating a true reduction of dimensionality requires showing that any combination of variables that leads to the same point in the low-dimensional space produces the same response. This is challenging, first because behavioral experiments tend to have a large variability which may hide small effects, and second because a convincing proof must test systematically a large number of equivalent combinations. Reaching at the same time high accuracy and a high number of combinations is beyond the experimental throughput in most behavioral experiments.

To address these challenges, we developed a high-throughput pipeline to study the foraging behavior of the nematode *Caenorhabditis elegans*. We focused on foraging (i.e. search and exploitation of food) because it has a clear impact on fitness, the degree of success is relatively easy to measure (in terms of rate of food consumption), and it is thoroughly studied from a theoretical point of view.^10^ Thanks to *C. elegans’* high offspring and small size, we could perform experiments with more than 20 000 age-synchronized individuals in more than 2 000 experimental arenas Besides allowing for high experimental throughput, *C. elegans’* small nervous system (∼300 neurons), makes it an ideal candidate to implement true low dimensional rules of thumb, while its foraging behavior is complex enough to implement the basic elements of optimal foraging, with well adapted behaviors for exploration^11–18^, learning^19–22^, and feeding^23–34^.

We systematically characterized *C. elegans’* response to food, covering all relevant combinations of food density (across 4 orders of magnitude, from starvation to rich environment) and food composition (across 12 different bacterial strains, from 11 diverse species). Different bacterial strains differ in their composition in terms of many different molecules, as well in their size, shape and mechanical characteristics, encompassing a high number of variables. Despite this high degree of complexity, our experiments revealed that *C. elegans* response to all bacterial strains follows a universal one-dimensional trend.

## Results

### *C. elegans’* reaction to food density follows a sigmoidal trend

Our experimental setup consisted of round agar plates with 5 food patches of different densities, arranged as a regular pentagon (**Figure 1a**). *C. elegans* is a bacteriophage, and each food patch was a drop of bacterial culture whose density had been carefully adjusted by measuring its optical density (OD). Approximately 10 young adult worms were placed at the center of the plate, equidistant to all food patches, and freely explored the environment for 2 hours, a time that was short enough to prevent significant food depletion, but long enough for patch occupancy to be roughly constant at the end of the experiment. To ensure repeatability and sufficient throughput, both the food patches and the worms were placed on the plate by a pipetting robot (see **Methods** for further details).

**Figure 1.**
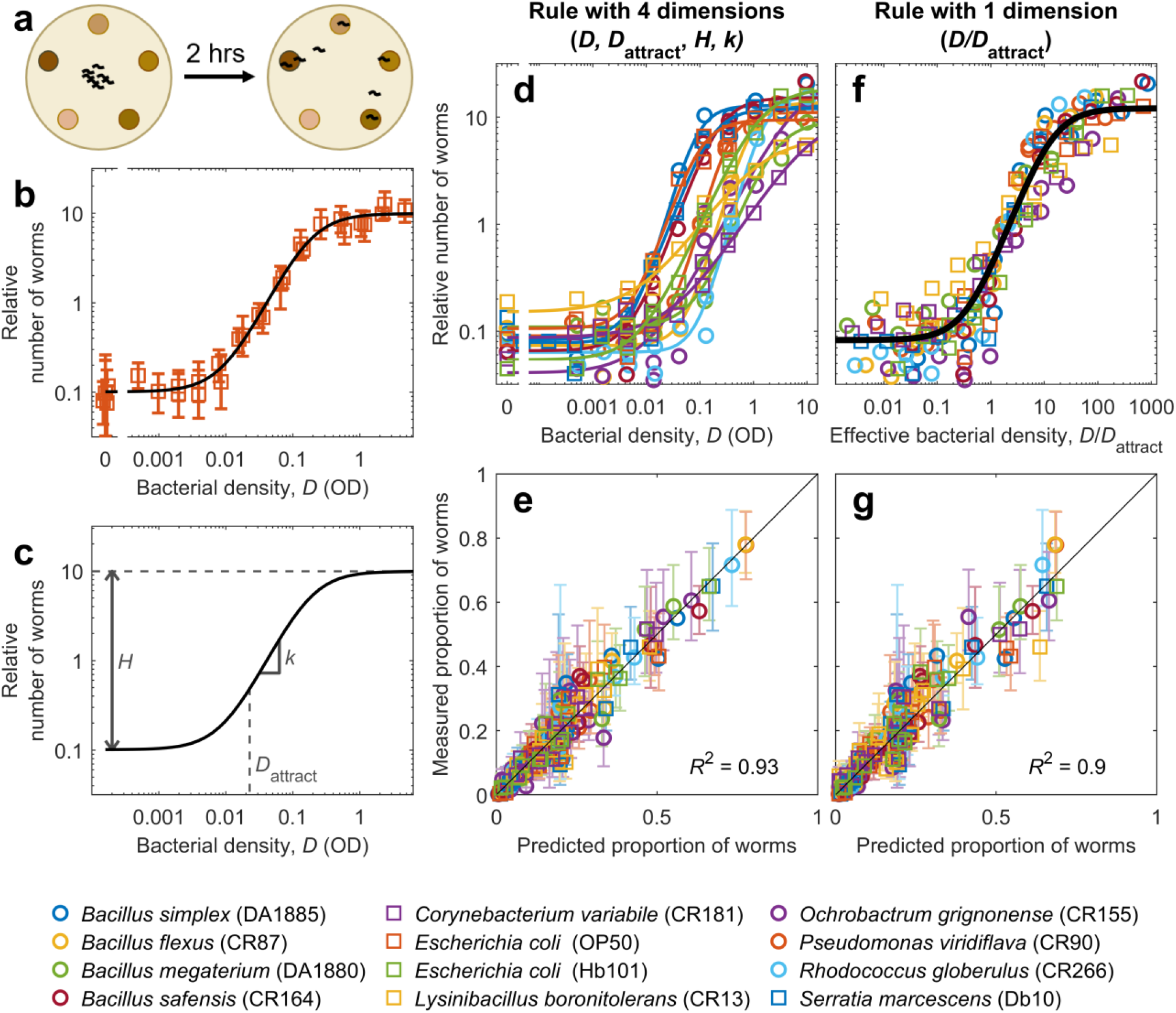
*C. elegans’* response to bacterial density follows a universal sigmoidal trend. **a**. Experimental scheme: Worms are placed at the center of a regular pentagon formed by 5 food patches of different densities. After 2 hours of exploration, worms located at each food patch are counted. **b**. Relative number of worms found at each food patch, as a function of bacterial density in the food patch (*D*). Squares: Experimental data for *E. coli* OP50; errorbars show the 95% confidence interval, computed via bootstrapping. Line: Fitted sigmoid, following **Equation 1 in Methods. c**. Sigmoid’s parameters: *H* is the ratio between the number of worms at the high and low density extremes, *k* is the slope at the sigmoid’s midpoint, and *D*_attract_ is the density at which the number of worms reaches 5-fold the low-density baseline. **d**. Same as (b), but for all bacterial strains and without errorbars (see **Figure S2** for separate plots for each strain and **Figure S3** for the parameters of all sigmoids) **e**. Measured proportion of worms in each food patch, versus proportion predicted by the sigmoid, fitted to each strains (**Equations 1** and **2**). Errorbars show the 95% confidence interval, computed via bootstrapping. **f**. Relative number of worms found at each food patch, as a function of effective density (*D/D*_attract_). Black line: Sigmoid with *H* = 146, *k* = 1.4. **g**. Same as (e), but with predictions made using effective density and the same sigmoid for all strains.

A key challenge was to study a wide enough range of bacterial densities, because placing patches of very different densities on the same plate leads to noisy data: Worms accumulate on the high-density patches, leaving very few individuals to assess the low-density range. We solved this challenge by performing several experiments covering smaller and overlapping density ranges. We then normalized the number of worms in each experiment with respect to a reference, to obtain a relative number of worms comparable across conditions (**Figure S1** and **Methods**).

We found that the relative number of worms at a food patch increases with its bacterial density following a sigmoidal trend, best visualized a double-logarithmic plot (**Figure 1b**). While our results are qualitatively comparable to previous studies^18,25,31,33^, our high throughput enabled a quantitative description and revealed that *C. elegans’* preference is well described by a mathematical formula with three parameters (**Figure 1c**; **Equation 1** in **Methods**): The height of the sigmoid (called *H* and defined as the ratio between the number of worms at the high and low density extremes), the speed at which the number of worms increases with bacterial density (called *k* and defined as the slope at the sigmoid’s midpoint), and the bacterial density at which the sigmoid starts to grow (called *D*_attract_ and defined as the density at which the number of worms reaches 5-fold the number of worms found at a control patch without bacteria). We can understand this last parameter as the minimum bacterial density needed to attract the worms significantly, so we called it “attraction density”. After fitting these three parameters, our sigmoid describes the experimental data remarkably well (**Figure 1b**, black line).

### *C. elegans’* response follows the same trend for all bacterial strains

To investigate whether our results are applicable to different types of food, we performed our experiments with 12 different strains, distributed across 11 species and 7 families. Five of these strains are common in *C. elegans* studies, while the remaining 7 strains are bacteria that we isolated from the gut of *C. elegans* (see **Methods**).

We found that the responses of *C. elegans* to all bacterial strains can be described by our sigmoidal equation, after fitting its three parameters independently for each strain (**Figure 1d** and **Figures S2, S3**). To quantify the overall goodness of this fit, we compared the proportion of worms at each food patch for each experimental condition with the predictions of our sigmoidal model (**Figure S1b**). We obtained an excellent agreement, with our model describing 93% of all experimental variance (**Figure 1e**).

We now turn to the dimensionality of the rule we found. We define dimensionality as the number of variables needed to describe how behavior changes in response to environmental change. Note that this definition is not necessarily the same as the total number of parameters of the model, because hard-wired parameters don’t count. In other words, we define the dimensionality as the number of parameters that need to be re-fit when the environment changes. The rationale of this definition is that it helps identify information bottlenecks: While our model is purely behavioral and does not intend to describe *C. elegans’* neural computations, the discovery of a low-dimensional rule that describes a wide range of experimental conditions would suggest an information bottleneck in *C. elegans’* nervous system.

According to this definition, our current model has dimensionality 4, since changing a food patch may lead to changes in any of the 4 parameters of our model: The density of the food patch (*D*), and the three parameters of the sigmoid (*H, k, D*_attract_), which may change from one bacterial strain to another. Therefore, while bacterial patches may differ in a myriad of parameters (their density, the size and shape of the bacteria, their hardness, their chemical composition, etc.), the impact of all these differences on patch occupancy must eventually reduce to changes in one or several of these four parameters. However, we asked if we could find an even simpler rule that still described our dataset with high accuracy.

### *C. elegans* responds to an effective bacterial density

The sigmoids shown in **Figure 1d** look remarkably similar, their biggest difference being a horizontal shift, which is controlled by the attraction density (*D*_attract_). Indeed, we found that differences across bacterial strains in parameters *H* and *k* are small, while *D*_attract_ differs up to 50-fold across strains (**Figure S3**).

We hypothesized that *C. elegans* might address the differences across bacterial strains simply by computing an effective density and reacting to it. We defined effective density as *D/D*_attract_, and found that this re-scaling of bacterial density removed most of the differences across bacterial strains (**Figure 1f**). We therefore re-fitted all our data with a single sigmoid, and found that we can describe all strains with the same parameters (*H* = 146, *k* = 1.4, black line in **Figure 1f**). This common sigmoid describes all our data almost as accurately as the separate fits (**Figure 1g**).

The rule describing *C. elegans* behavior is thus reduced to one dimension, since now the sigmoid’s parameters *H* and *k* do not depend on the environment (i.e. they can be hard-wired), and the effective density *D/D*_attract_ is the only variable that needs to change when the environment changes.

### Preference correlates with fitness gained from each strain

We then asked whether *C. elegans’* response was well adapted to choose the food patches that maximize its fitness. We used offspring as a proxy of fitness, counting the number of eggs laid by an individual worm that feeds on a food patch for 5 hours. In order to remove the effect of food preference as much as possible, we encircled the food patch with a copper ring that prevented the worm from escaping, forcing it to stay on the food patch regardless of its preference (**Figure 2a**). We measured this proxy of fitness for every bacterial strain and over a wide range of densities, and found that it increases sharply at a given food density that we called *D*_fitness_, and stabilizes at higher densities (**Figure 2b**).

**Figure 2.**
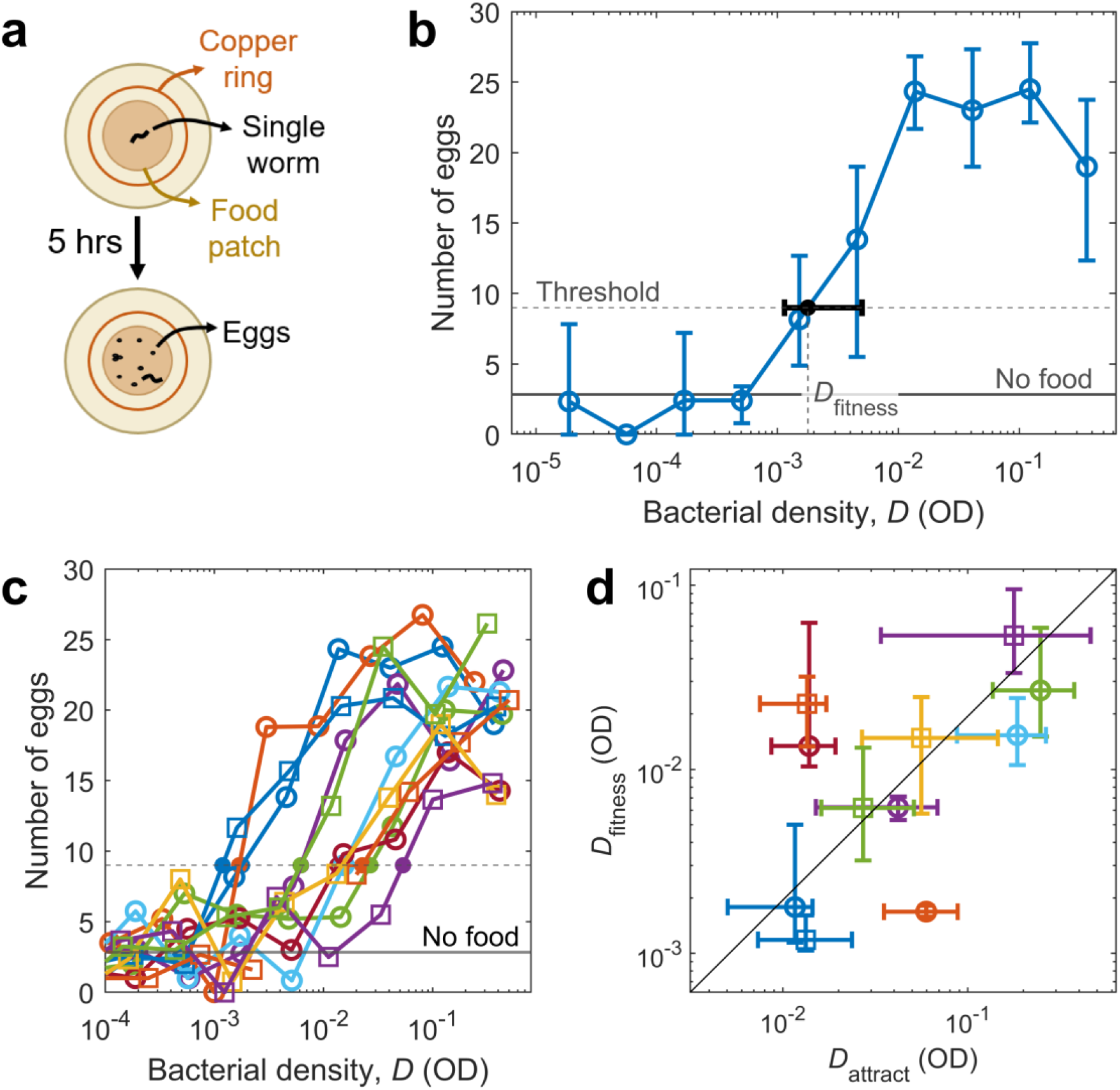
Attraction to each strain correlates with fitness benefit. **a**. Experimental scheme to estimate fitness: A single worm was placed in the middle of a food patch, and surrounded by a copper ring to prevent it from escaping. Five hours later, the number of eggs was counted. **b**. Number of eggs laid after 5 hours on a food patch of DA1885, as a function of the density of the food patch (open circles). Solid horizontal line: Average number of eggs when no food was present. Dashed horizontal line: Threshold chosen to determine when the number of eggs increases with respect to the no-food baseline. Solid dot: Point at which the number of eggs crosses the threshold, which defines *D*_fitness_. **c**. Same as (b), but for 11 bacterial strains and without errorbars. **d**. Density at which worms start laying more eggs (*D*_fitness_), versus density at which worms start being attracted to a food patch (*D*_attract_). Line indicates perfect proportionality between the two variables (*D*_fitness_ = 0.2*D*_attract_; the value of the proportionality constant has little consequence, since it depends on the thresholds chosen to define *D*_attract_ and *D*_fitness_). Color and shape of all markers identify bacterial strain, following the legend in **Figure 1**. All errorbars show 95% confidence intervals, calculated via bootstrapping.

Fitness increases with bacterial density following a similar trend for all strains, the main difference being a shift in the density at which the increase takes place (*D*_fitness_) (**Figure 2c**). Therefore, an optimal behavioral response should also follow the same trend for all strains with a shift in density, which is what we found for patch occupancy (**Figure 1**). The question is whether the density shifts in food preference (characterized by the attraction density *D*_attract_) correspond to those found in fitness (characterized by *D*_fitness_). We found this to be the case: *D*_fitness_ and *D*_attract_ are proportional (*p* = 0.03, linear regression), which in a log-log plot corresponds to a line with slope 1 (black line in **Figure 2d**).

Three outliers deviate from the general trend (**Figure 2d**). One of these outliers is *E. coli* OP50, which was also used to feed the worms before the experiment. This previous experience might explain the deviation, as it might increase the worms’ preference for OP50 with respect to unfamiliar strains^19,21^. The other two outliers (*Bacillus safensis* CR164 and *Pseudomonas viridiflava* CR90) cannot be explained in this way, and are probably due to factors that impact fitness but are neglected by the worms. These deviations suggest that *C. elegans’* behavior is near-optimal but not perfectly optimal, although we must keep in mind that we only measure a proxy for fitness (number of eggs laid in 5 hours), and a more accurate measurement of fitness might partially explain the outliers. In any case, we conclude that *C. elegans* is using a rule of thumb, focusing on cues that allow it to adapt its behavior to most strains, and probably neglecting others that would be relevant for the outliers.

### Biomass content does not drive food choice

We next asked what sensory cues determine the observed rule of thumb. The amount of food eaten should be the main driver of foraging behavior, and the amount of food actually available for the worms might be different for different strains, even at the same OD. OD measures the amount of light absorbed by a bacterial culture and is proportional to biomass density for a given bacterial strain. However, different bacterial strains have different cell size, shape and composition, which affect light transmission through the culture. Therefore, cultures of different strains at the same OD may have different biomass density. We hypothesized that the different values of attraction density (*D*_attract_) in terms of OD might in fact reflect the same density threshold in terms of biomass.

To measure biomass density of each bacterial strain, we determined the relation between biomass content and OD. We did this by measuring the weight of dry biomass left after evaporating all the water contained in bacterial cultures of all our strains. If biomass differences were to explain our results, we should find an inverse relationship between biomass content and *D*_attract_, because strains with twice as much biomass at OD=1 should have an effective density twice as large, and therefore their attraction density (*D*_attract_) should be halved.

While we did find a slight negative correlation (*p*=0.05, linear regression) between biomass content and *D*_attract_, this correlation is far too weak to explain the differences across bacterial strains: Biomass density changes by less than 3-fold across different strains, while *D*_attract_ changes up to 50-fold (**Figure 3a**; if differences in biomass density were to explain differences in *D*_attract_, the datapoints should follow the black diagonal, which has slope −1 in this log-log plot). Therefore, biomass density is not the main driver of food choice in *C. elegans*.

**Figure 3:**
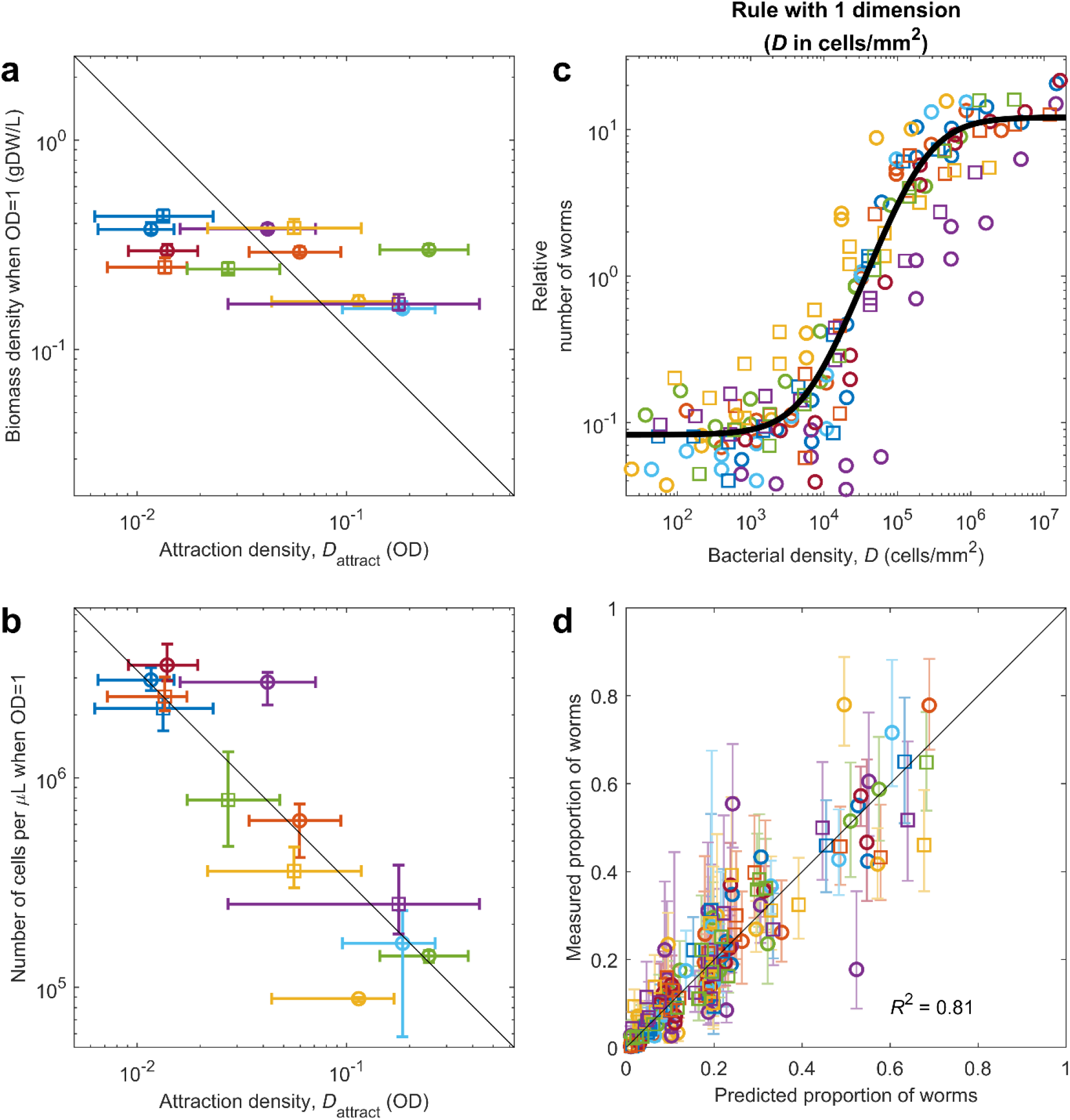
*C. elegans’* rule of thumb is driven by the number of bacteria per unit surface. Color and shape of markers identify bacterial strains (see legend in **Figure 1**). **a**. Biomass density for each strain at OD=1 (measured in grams of dry weight, gDW, per liter), versus its attraction density (*D*_attract_). Black line: Inverse relation (proportional to 1/*D*_attract_), which would indicate that biomass density is responsible for the observed differences in *D*_attract_. **b**. Number of cells per microliter at OD=1 for each strain, versus *D*_attract_ for each strain. Black line: Inverse relation (proportional to 1/*D*_attract_), which would indicate that the number of cells is responsible for the observed differences in *D*_attract_. **c**. Relative number of worms found at each food patch, as a function of bacterial density (measured in cells/mm^2^) in the food patch. Black line: Sigmoid, fitted to all strains. **d**. Measured proportion of worms in each food patch, versus proportion predicted by the sigmoid in (c). Errorbars show the 95% confidence interval, computed via bootstrapping.

### Cell density drives food choice

Next, we hypothesized that cell density might be driving the preference. For the same reasons discussed in the previous section, different bacterial strains will have different cell density (i.e. number of cells per unit volume) at the same OD. We determined the cell density in our cultures by a combination of plating and microscopic observations (see **Methods**). As for biomass, we expected to find an inverse relation between *D*_attract_ and cell density at OD=1.

We found an excellent inverse correlation between *D*_attract_ and cell density at OD=1 (*p*=0.002, linear regression), and in this case the correlation was strong enough to explain all the variability in *D*_attract_ (**Figure 3b**). This result indicates that the effective density that drives *C. elegans* behavior is simply bacterial density, but measured in number of cells per unit of volume (or number of cells per unit of surface, once the bacteria are placed on the surface of the agar plate). We confirmed this fact by plotting our original data with the bacterial density measured in cells/mm^2^, and finding that all sigmoids collapse to a great extent (**Figure 3c**).

Therefore, a one-dimensional rule that characterizes a food patch exclusively by its density in cells/mm^2^ (making no distinction across bacterial strains), describes the experimental results with high accuracy (**Figure 3d**). However, we do find a slight drop in accuracy: Our previous model explained 90% of all experimental variance (*R*^2^ = 0.9, **Figure 1g**), while the current one explains 80% of it (*R*^2^ = 0.8, **Figure 3d**). This drop in accuracy probably comes from a combination of two issues: First, other factors besides bacterial cell density may contribute to the effective density that drives *C. elegans*’ behavior, so this drop in accuracy may reflect actual biological complexity. Second, our current model has substituted the *D*_attract_ that were fit to our behavioral results for an independent measurement of the cell density of each bacterial strain, and the experimental inaccuracies of this separate measurement must necessarily reduce the accuracy of the overall fit.

In any case, our results robustly indicate that at least 90 % of *C. elegans’* response to bacteria is driven by a one-dimensional rule of thumb (**Figure 1g**), and at least 80 % of the response can be explained by a single environmental variable (bacterial cell density, **Figure 3d**), which is the most informative one in terms of fitness benefit (as compared to alternatives, such as biomass density).

### Response to mixed food patches confirm our results

Our results indicate that *C. elegans* simply measures the number of bacteria encountered per unit surface when deciding whether to keep exploiting a food patch or to leave it. This result produces an interesting prediction, which in turn provides a stronger test of our hypotheses. Let’s consider a food patch in which two bacterial strains are well mixed. A worm exploring this food patch will be simultaneously exposed to both strains, so any factors such as different cell composition, different cell size or different metabolites, will be perceived near-simultaneously. We don’t know how these stimuli combine in *C. elegans’* nervous system, so if they play an important role it would be hard to predict *C. elegans*’ response to a mixed food patch. As a reference we define a reasonable null model, assuming that the response to a mixed food patch would be an average of the responses to the corresponding pure patches, weighted by their relative proportions in the mixture. Therefore, we define our null model as any result between the weighted arithmetic and geometric means of the responses to each strain separately (**Figure 4a**).

**Figure 4:**
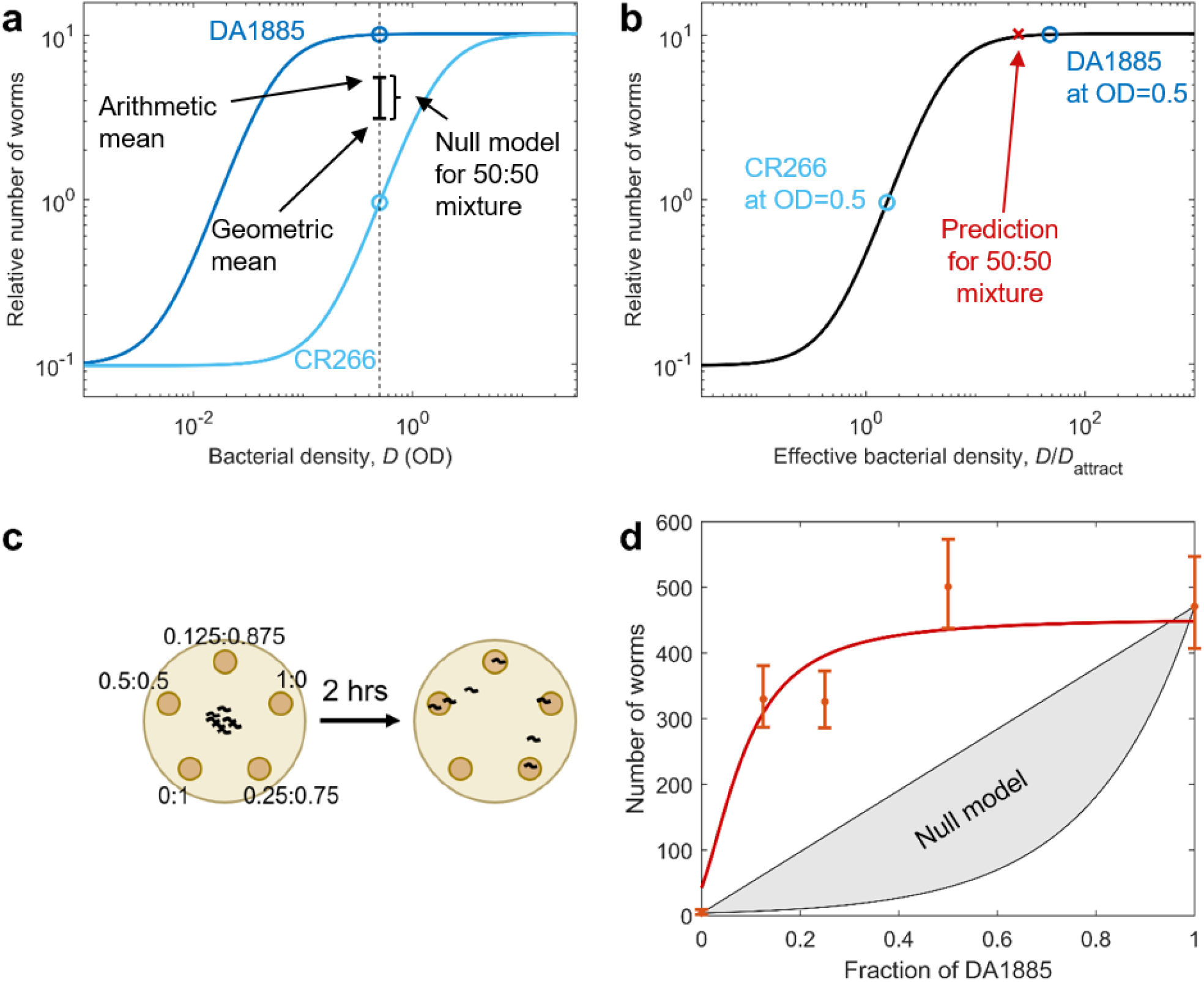
The response to mixed food patches shows that effective density is additive. **(a)** Relative number of worms predicted by the sigmoidal model for patches of DA1885 (dark blue) and CR266 (light blue), as a function of bacterial density (measured in OD). Our experiment took place at OD=0.5 (black dashed line), where DA1885 is about 10 times more attractive than CR266 (circles). The null model for a 50:50 mixture of both strains goes from the geometric mean to the arithmetic mean of the two pure patches (black errorbar). **(b)** Relative number of worms in a food patch, as a function of its effective bacterial density (*D/D*_attract_). Circles: Effective density for CR266 and DA1885 at OD=0.5. Red cross: Effective bacterial density for a 50:50 mixture of CR266 and DA1885 at OD=0.5 (the effective density of the 50:50 mixture is the arithmetic mean of both effective densities, which in a logarithmic scale is located closer to the highest one). **(C)** Experimental scheme: Worms were exposed to 5 food patches with different fractions of CR266 and 1885, always at OD=0.5. Worms at each patch were counted after 2 hours. **(D)** Number of worms at each food patch, as a fraction of DA1885. Dots: Experimental results (errorbars show the 95% confidence interval). Red line: Prediction from the sigmoidal model (as in box b). Gray patch: Prediction of the null model.

In contrast, if *C. elegans’* response is determined by the cell density of the food patch, we can predict the response to any mixed patch, by computing its effective density (which is the average of both strains’ effective densities, because cell density is additive), and use our sigmoidal model to predict the worm’s response to it (**Figure 4b**). By choosing pairs of bacterial strains with very different *D*_attract_, we can have cases in which this predicted response is very different from the null model (compare **Figure 4a** and **Figure 4b**).

To test these predictions, we performed experiments with food patches containing mixtures of CR266, which has a high *D*_attract_, and DA1885, which has a low *D*_attract_. We prepared cultures at OD=0.5 for both strains, and prepared 5 mixtures with ratios DA1885:CR266 of 0:1, 0.125:0.875, 0.25:0.75, 0.5:0.5, and 1:0. Given that both pure cultures had OD=0.5, all 5 mixtures also had OD=0.5 (range 0.49 to 0.51). We then run our experiment, letting worms choose among 5 food patches made from these 5 mixtures (**Figure 4c**).

The experimental results follow the predictions of the sigmoidal model, and clearly reject the null model (**Figure 4d**). Besides the pair of strains presented here, we measured another three pairs (a total of four). Three out of the four pairs showed excellent agreement with our predictions, and in all cases cases we found better agreement with our predictions than with the null model (**Figure S4**).

## Discussion

Our results show that *C. elegans’* response to food across bacterial species is driven by a single variable: Effective food density. This variable seems to correspond to bacterial cell density, with other factors that depend on bacterial strain, such as biomass content, having minimal effect.

Our experimental results may seem to contradict previous studies, but are consistent with them. Previous studies^25,26^ showed large differences in preferences between certain strains, such as *E. coli* Hb101 and *E. coli* DA837, which is very similar to OP50 and elicits the same behavioral response (**Figure S5**). In contrast, we found only moderate differences between Hb101 and OP50. But studies showing larger differences were performed on nutrient agar plates, so bacteria could grow after being deposited on the plate. OP50 is a uracil auxotroph, and this fact limits its growth on solid media, while Hb101 does not have this limitation and grows to higher densities on agar plates. Therefore, differences across strains observed in previous studies may be attributed to differences in bacterial density. Another apparent contradiction is the evidence that less-preferred bacteria are hard to eat for *C. elegans*^25,35^. We would not expect that increasing the density of hard-to-eat bacteria would make them as profitable as easier-to-eat strains, so the regularity of our results challenges this previous finding. However, the main reason why some bacterial strains were hypothesized to be harder to eat was that they had a larger cell size,^25,35^ and strains with larger cell sizes tend to have smaller cell densities at saturation. Therefore, the differences observed in previous studies were also correlated with cell density. A final apparent contradiction comes from studies that have shown that *S. marcescens* (Db10) is a pathogen of *C. elegans*, and actively avoided by the worms^36,37^. We did not find such avoidance behavior, but both the strong pathogenicity and the avoidance response require active production of a toxin by the bacteria, which was not possible in our experimental conditions due to the lack of nutrients in the plates. As a control, we checked that we could reproduce *C. elegans’* avoidance of *S. marcescens* when performing experiments on NGM plates rather than on our experimental plates (**Figure S6**). In sum, revealing *C. elegans’* foraging rule of thumb required accurate control of bacterial density and decoupling the effect of toxins and other metabolites.

A limitation of our study is that we measured a single experimental outcome (patch occupancy). Differences in this outcome emerge from changes in elementary behavioral parameters, such as speed, turning rate, probability of different behavioral states (such as roaming, dwelling and quiescence), etc.^23,24,27,31^ A more detailed study of these parameters might reveal differences that were not apparent here. This higher degree of detail was not possible at the level of throughput and coverage needed to reveal the rule of thumb, but is a natural next step towards unveiling its mechanistic and neural implementation.

A second limitation of this study is the lack of bacterial growth, which limits the production of bacterial metabolites. These metabolites are probably relevant in natural conditions, as is certainly the case for pathogenic bacteria^36,37^. Our experimental conditions were necessary to properly control bacterial density, and the good correlation of behavior and fitness benefit shows the ecological relevance of our observations. These controlled experimental conditions have revealed a core behavioral mechanism, which produces adaptive response by focusing on the single-most informative environmental variable.

## Materials and Methods

### Strains and media

All experiments were performed with *Caenorhabditis elegans*, strain N2, obtained from the Caenorhabditis Genetics Center (University of Minnesota, https://cgc.umn.edu/), and maintained using standard practices.^38^ Worms grew at 22 C on nematode growth medium (NGM: 3 g/L NaCl, 2.5 g/L peptone, 20 g/L agar, 25 mL/L potassium phosphate buffer pH 6, 1 mM MgSO_4_, 5 mg/L cholesterol, 1 mM CaCl_2_), in 100mm Petri dishes seeded with 200μL of a saturated culture of *E. coli* OP50 bacteria (grown overnight on LB at 22 C). The worms were transferred to a fresh dish every 1 to 3 days to prevent food depletion, so that worms used in any experiment came from a population that had not experienced food depletion for at least 5 generations. To prevent accumulation of mutations, we ensured that our population was never more than 30 generations away from the individuals received from the CGC. We performed all experiments with 48-old worms, synchronized by bleaching and egg collection.

*Escherichia coli* (OP50), *Escherichia coli* (Hb101), *Escherichia coli* (DA837), *Serratia marcescens* (Db10), *Bacillus megaterium* (DA1880), and *Bacillus simplex* (DA1885) were obtained from the Caenorhabditis Genetics Center. *Lysinibacillus boronitolerans* (CR13), *Bacillus flexus* (CR87), *Pseudomonas viridiflava* (CR90), *Ochrobactrum grignonense* (CR155), *Bacillus safensis* (CR164*), Corynebacterium variabile* (CR181), *Rhodococcus globerulus* (CR266), *Pseudomonas veronii* (CR220), and *Raoultella terrigena* (CR225) were isolated by us from the gut of *C. elegans* N2 worms who had fed on organic compost (see below).

Bacteria were streaked on NGM plates from a −80 C glycerol stock, stored at 4 C, and re-streaked to a fresh plate every two weeks to ensure viability. To prepare liquid cultures, we inoculated one or two bacterial colonies in 5mL of LB medium, and incubated for 24 h, at 22 C, with orbital shaking at 300 rpm, in a closed 50mL Falcon tube. Then, 1μL of this culture was inoculated in either 5 or 10mL of fresh LB and incubated for another 24 h in the same conditions. *E. coli* Hb101 was an exception, as it took longer than 24 hours to reach saturation. In this case we skipped the second inoculation, continuing the incubation of the original culture for a total of 48 hours.

### Isolation of *C. elegans* gut bacteria

The natural microbiota strains of *C. elegans* were isolated by growing *C. elegans* on different types of rotten organic material, followed by washing and sterilizing the worms on the outside, grinding the worms and plating the resulting bacterial suspension on agar plates.

We first prepared heat killed *E. coli* OP50 by growing OP50 for 24h in 200mL tryptic soy broth (Teknova, Hollister, CA, USA) at 37°C, followed by spinning down, resuspending in 4mL S-medium (prepared as described in ^38^) and incubation at 80°C for 24h. This procedure results in 50x *E. coli* OP50, 50x compared to density at saturation

Two types of food sources were fed to the worms: different types of (i) compost and (ii) rotten fruits and vegetables. Some rotten apples were directly collected from the outside. Other fruits like apples, celery, almonds and parsnip were placed on local soil from Boston, MA in a household plastic box (Sterilite) with semi-open lid and incubated in the lab at room temperature until the fruits were strongly decayed (∼3 weeks). The compost samples were taken from two local compost piles in Boston, MA, that mostly contained kitchen and garden waste. Some amount of phosphate buffered saline and glass beads were added to the samples. The samples were homogenized by vortexing at high speeds. The resulting solution was filtered through a 5µm filter (Millex-SV 5.0 µm, MerckMillipore, Darmstadt, Germany) to remove bigger particles. The resulting emulsion was spread on S-media agar plates without citrate.

*C. elegans* N2 were first grow on *E. coli* OP50 lawn on NGM plates. The worms were washed off the plates with M9 worm buffer with 0.1% Triton X-100. The worms were let sink down for about 1min and the supernatant was removed. The worms were resuspended in S-medium containing 100µg/mL gentamicin and 5x heat-killed OP50 (5x compared to density upon saturation). The worms were incubated in that solution for 24h at room temperature with gentle shaking (50mL tube, semi-unscrewed cap). Finally, the worms were washed twice with M9 worm buffer + 0.1% Triton X-100.

The germ-free worms were added to the plates with rotten organic material for around one week. After that time the worms were washed off the plates with M9 worm buffer with 0.1% Triton X-100. The worms were washed twice with M9 worm buffer with 0.1% Triton X-100 (centrifugation at 2000g, 10s). Afterwards the worms were re-suspended in 1mL ice cold M9 worm buffer with 0.1% Triton X-100 and incubated on ice for 10mins. 2µL bleach (Clorox) were added to kill bacteria on the outside of the worm and the worms were incubated for 6mins on ice. Afterwards the worms were washed 3x with ice cold M9 worm buffer with 0.1% Triton X-100. Single worms were transferred into 0.6mL reaction tubes (Eppendorf) and ground with a motorized pestle (Kimble Kontes Pellet Pestle, Fisher Scientific) for at least 1min. The resulting solution was plated onto a tryptic soy broth (Teknova, Hollister, CA, USA) agar plate (2% agar, 150mm petri dish). From the resulting colonies, physiologically unique colonies were picked. The colonies were streaked out again on tryptic soy broth agar and checked for contaminations. If contaminations were spotted the bacteria were re-streaked again. Finally, the bacteria were grown in tryptic soy broth at 30°C and stored as glycerol stocks. The species identity was analyzed by 16S Sanger sequencing (Genewiz, South Plainfield, NJ).

### Preparation of experimental plates

Assays were run in foraging plates (3 g/L NaCl, 20 g/L agar, 25 mL/L potassium phosphate buffer pH 6, 1 mM MgSO_4_, 5 mg/L cholesterol, 1 mM CaCl_2_, 10 mg/L chloramphenicol and 100 mg/L novobiocin). The composition of these plates was designed to prevent bacterial growth, not containing any nutrients for the bacteria, and containing two bacteriostatic antibiotics. We chose this antibiotic cocktail after measuring the Minimum Inhibitory Concentration (MIC) for 6 different bacteriostatic antibiotics and all our bacterial strains. We aimed to prevent bacterial growth while keeping the bacteria as healthy as possible, and we determined that 10 mg/L chloramphenicol and 100 mg/L novobiocin was the best combination to prevent the growth of all strains while keeping the antibiotic concentrations as low as possible. We checked that all bacterial strains remained viable and with constant optical density after 24 hours of exposure to this cocktail of antibiotics. Plates were poured one week before the assays, and stored at room temperature.

One day before the experiment, bacterial cultures were washed three times with foraging buffer (3 g/L NaCl, 20 g/L agar, 25 mL/L potassium phosphate buffer pH 6, 1 mM MgSO_4_, 1 mM CaCl_2_, 10 mg/L chloramphenicol and 100 mg/L novobiocin). After the last wash, we re-suspended the bacteria in foraging buffer, adjusting their OD with a spectrophotometer (Jenway 7200, Cole-Parmer, Staffordshire, UK) to the maximum OD needed for our experiment. We then performed serial dilutions in foraging buffer to obtain all needed densities.

A pipetting robot (OT-2, Opentrons, Long Island City, NY, USA, with custom modifications to handle agar plates) placed drops of bacterial culture on the foraging plates. In all cases, we used 40 μL drops, which spread to a diameter of 11.2 mm on average. Drops were left to dry overnight at 22 C.

### Patch occupancy assays

We used 55mm-diameter foraging plates with five 40-μL drops of bacteria, forming a regular pentagon with the patch centers at 13 mm from the plate’s center. For each experimental condition (consisting of 5 different food densities), we prepared at least 4 different versions, randomly permuting the position of the 5 densities across the 5 food patches to minimize effects due to relative position of the food patches. We then prepared at least 8 replicates of each version, so we had a total of at least 32 plates per condition. We also randomized the order at which the different conditions were prepared. Food patches were placed on the experimental plates one day before the experiment and dried overnight at 22 C.

48-hour old synchronized worms were washed off their cultivation NGM plates with M9 worm buffer + 0.1% Triton X-100 (3 g/L KH_2_PO_4_, 7.52 g/L Na_2_HPO_4_.2H_2_O, 5 g/L NaCl, 1 mM MgSO4, 0.1% Triton X-100; Triton X-100 was added to prevent worms from sticking to the pipette tips). To remove all bacteria, we washed the resulting worm suspension 6 times with M9 + 0.1% Triton X-100 using a table-top centrifuge (∼5 second spin was enough to pellet the worms while leaving the bacteria in suspension). Worms were then placed in the middle of the experimental plates by the pipetting robot, in drops of 10-15μL (adjusted for an average of 10 worms per plate). The full wash procedure took between 8 and 10 minutes and placing the worms took at most 5 minutes, so at most 15 minutes elapsed between the breeding plate and the experimental plate.

The worms were left on the plates for 2 h at 22 C, and then we imaged the plates at 1200 dpi using a scanner (Epson Perfection V850 Pro). Previous protocols that use similar scanners to quantify worm behavior proposed modifications to increase image quality and control temperature.^39^ However, we found that unmodified scanners provided good enough image quality for our purposes, and temperature control was not an issue for us because plates were placed on the scanners only briefly at the end of the experiment. Extreme care must be exercised when placing the plates on the scanners, since even a gentle tap may startle the worms and make them leave the food patches. Worms were automatically located in the images using a custom-made program built in Matlab R2019a.

We aimed to having 32 plates for each experimental condition (8 plates for each version with permuted positions), with 10 worms per plate. However, the actual number of plates and worms was lower. First, we removed a small fraction of plates that presented imperfections on the agar surface or non-round food patches. Second, given that we could not control the exact number of worms placed on each experimental plate (just the volume and concentration of worm suspension), we had substantial variability in worm number per plate. To prevent any significant effects from food depletion, we removed from the analysis all plates that had more than 20 worms. After this filtering, we had 29 ± 4 plates per condition and 230 ± 110 worms per condition (mean ± standard deviation). Then, for each experimental condition, we added up all the worms found at patches of a given density (across all replicates), and added one pseudocount to obtain a less biased estimate.^40^

### Fitting and normalization of patch occupancy data

We assume that the number of worms found in a given food patch is proportional to its attractiveness, *A*, which we define as

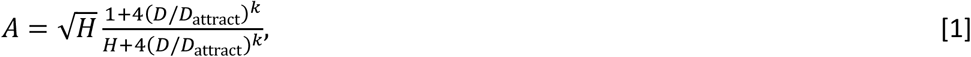

where *H* is the ratio between its highest and lowest points, *k* is the slope at the sigmoid’s midpoint (in a double-logarithmic plot), and *D*_attract_ is the density at which the relative number of worms reaches 5-fold the low-density baseline (**Figure 1c**; the low-density baseline is 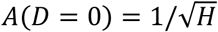, and 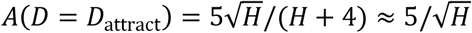 when *H* ≫ 1).

Then, the proportion of worms present in each food patch in a given plate will be

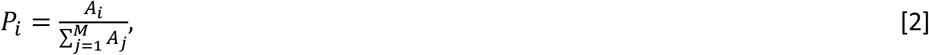

where *P*_*i*_ is the proportion of worms in the *i*-th food patch, *A*_*i*_ is the attractiveness of the *i*-th food patch, and *M* is the number of patches present in the experiment. **Equation 2** is a strong assumption, which holds approximately in our system for reasons that will be explored in a separate article, and which is validated by the excellent goodness of fit of our model (**Figure 1e**).

To fit the model’s parameters to our experimental data, we maximize the log-likelihood of the model. For one experimental plate, the log-likelihood is

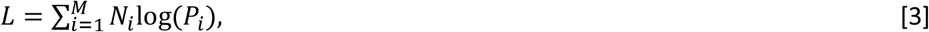

where *N*_*i*_ is the number of worms found in the i-th patch, *M* is the number of patches (*M*=5 in all our experiments), and *P*_*i*_is computed using **Equations 1 and 2**. We then added up the log-likelihoods of all experiments that we wanted to fit with the same set of parameters, and found the set of parameters that maximized this accumulated log-likelihood, using Matlab’s ‘fmincon’ function (Matlab R2019a). Once the optimal parameters are found, **Equations 1 and 2** provide a good description for the proportion of worms reaching each patch in each separate experiment (**Figure S1A**). These two equations are also used to represent all the “predicted vs experimental” plots presented in the paper (**Figures 1e, 1g, 3d**).

In order to show together the experimental data coming from experiments that cover different density ranges for the same bacterial strain (the three columns in **Figure S1**), we re-normalized the data and computed a relative number of worms, *N*_*i*_*/N*_ref_ (where *N*_*i*_ is the number of worms in the i-th food patch, and *N*_ref_ is the number of worms in a reference patch). This normalization would be trivial if we had a reference food patch of some common density in every experiment, but this was not possible in practice: Our range of densities is very wide, and differences in patch occupancy span more than 2 orders of magnitude. When food patches of very dissimilar densities are placed in the same plate, worms accumulate in the high-density ones leaving the low-density ones almost completely empty, and leading to very noisy datasets. For this reason, each individual experiment covered a relatively small range of densities (**Figure S1**). Therefore, while neighboring ranges overlap with at least two datapoints, we did not have any one density present in all of them. To circumvent this issue and be able to represent all the data together, we used our sigmoidal fit to estimate the number of worms we would expect at a virtual reference patch, and used this estimate to re-normalize our experimental data. We chose our virtual reference patch to be in the sigmoid’s midpoint, which led to a normalization factor

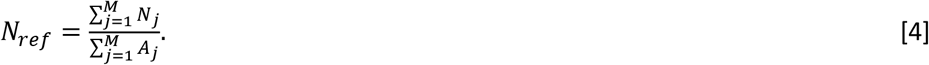

Due to this normalization, the relative number of worms is 1 at the sigmoid’s midpoint (see for example **Figure 1**).

This procedure re-aligns the experimental data from different experiments (**Figure S1c**), allowing us to present it in a single graph (**Figure S1d**). However, this alignment depends on the fit itself, and therefore does not provide a reliable visual indication of the goodness of the fit. This visual indication of the goodness of fit is found in the “predicted vs experimental” plots, which are not affected by this normalization (**Figures 1e, 1g, 3d**).

### Fitness experiments

We used 35mm foraging plates with one 40μL drop of bacteria in the center, placed on the plate the day before the experiment and dried at 22 C. 48-hour old worms were washed from their breeding plates in the same way as for patch occupancy assays. Then, individual worms were fished using a pipette and placed on the bacterial patch (one worm per plate). A copper ring of 2 cm diameter was then lodged into the agar, around the patch, to prevent the worm from escaping. Worms were left on the lawn for 5 hrs and then put at −20°C for 5 min. This brief period ensured quick refrigeration of the plates to immobilize the worms and stop egg-laying, without freezing the agar. Then, plates were stored at 4°C. Worms remained immobile and eggs didn’t hatch, so eggs could be counted for at least 2 weeks after the experiment. Eggs were manually counted, excluding any plates where more than one worm was placed by mistake, or where the worm escaped the area delimited by the copper ring. All experiments were performed on the same day to minimize experimental variability, but eggs were counted over the 2 weeks following the experiment. Because of experimental complications, we failed to measure *C. elegans* fitness on *B. flexus* (CR87), so we have measurements for 11 out of our 12 strains.

To determine *D*_fitness_, we first found the highest density for which the average number of eggs was below the threshold. Then, we performed linear interpolation between that point and the next one, with bacterial densities in logarithmic scale.

### Determination of bacterial density

Optical Density (OD) was measured using a spectrophotometer (Jenway 7200, Cole-Parmer, Staffordshire, UK). We found that this spectrophotometer is most accurate for OD’s between 0.1 and 1, so we always diluted the bacterial cultures to obtain measurements in this range. Lower OD’s could not be measured accurately, so they are inferred from the dilution factors used to prepare them.

To determine density in cells per microliter, we combined plating to determine the amount of colony forming units (CFU) with microscopy to investigate the nature of each CFU.

We determined the CFU density as follows. After determining the OD of the bacterial culture, we performed 10-fold serial dilutions in M9 worm buffer (3 g/L KH_2_PO_4_, 7.52 g/L Na_2_HPO_4_.2H_2_O, 5 g/L NaCl, 1 mM MgSO4), and plated four 10-microliter drops of each dilution on an NGM plate. We incubated this plate at room temperature for 48 hours, counted the number of colonies in the drops that had around 10 colonies, and used these counts to derive the density of colony-forming units in our original culture. We computed the errorbars by bootstrapping the four drops for each measurement. We performed this plating procedure both before and after washing the bacteria with foraging buffer to prepare our experimental plates (see “Preparation of experimental plates”), and we did not find any consistent differences before and after the wash.

The number of CFU/μL is not identical to the number of cells/μL. First, not all cells that fall on the surface of an agar plate survive and manage to form a colony. To control for this effect, we performed a control using LB plates instead of NGM plates, and we did not find significant differences in viability across these two types of plates, which suggests that viability was high for all strains in both media. Second, each CFU may be a single cell, but it may also be a cluster of cells that clump together and form a single colony. To control for this effect, we studied our bacterial cultures under a microscope (LEICA DM6000). We found that 10 out of our 12 bacterial strains were mostly composed of individual cells, with few clumps, so for these 10 strains CFU/μL is a good estimate of cells/μL. In contrast, *B. megaterium* (DA1880) and *B. flexus* (CR87), form long filaments composed of several cells, and each of these filaments will form a single colony. Using DAPI staining to visualize individual cells in each filament, we counted the number of individual cells per filament, obtaining 9.6 and 8.7 cells per filament on average for *B. megaterium* (DA1880) and *B. flexus* (CR87), respectively. We used these factors to transform the CFU/μL determined from plating to cells/μL for these two strains.

To determine the amount of biomass present in our bacterial cultures, we prepared 200 mL of saturated culture for all strains, washed it 3 times with M9 worm buffer, and resuspended to a volume of 5 mL. We then measured the OD of these suspensions, and placed them in glass tubes that we had previously weighed. We also added three tubes with M9 worm buffer without bacteria, to be able to account for the weight of the salts contained in the buffer. We evaporated all the water by incubating the tubes at 90 C for 24 hours, and weighed them again. We checked that longer incubation did not change the weight, meaning that 24 hours were enough for all the water to evaporate. We then calculated the biomass contained in each tube by subtracting the weight after incubation minus the weight of the empty tube, and minus the weight corresponding to the salts from the M9 buffer (to calculate this weight we followed the same procedure with tubes that contained only M9 buffer, obtaining a dry weight of 15 g/L, which is close to the theoretical weight we would extract from the recipe of M9 worm buffer). See detailed protocol at dx.doi.org/10.17504/protocols.io.kxygxzn44v8j/v1. To compute confidence intervals, we assumed that all weight measurements had the same proportional error observed in the 3 measurements performed to estimate the weight of M9 salts, we estimated the error in OD measurements by performing 10 measurements of the same culture, and we combined these two sources of error using bootstrap.

### Errorbars and statistics

We computed all errorbars using bootstrap^41^: For a given experimental condition for which we have *P* replicates (i.e. *P* experimental plates), we chose *P* of these replicates randomly, with replacement (so some replicates can be chosen several times, and some will not be chosen). By doing this with all of our experimental conditions, we obtained a bootstrapped dataset. We thus generated at least 1000 bootstrapped datasets. These bootstrapped datasets are an estimate of what we should expect if we repeated our whole experimental process 1000 times, so they give an estimate of the reproducibility of our results.^41^ For each errorbar shown in the paper, we computed the corresponding quantity for each of the bootstrapped datasets, removed the most extreme 2.5% of values at each side, and reported the remaining interval as an errorbar. For example, errorbars in *D*_attract_ were computed fitting our sigmoid to each of the bootstrapped datasets removing the most extreme 2.5% values of *D*_attract_ resulting from these fits, and reporting the remaining interval.

Significance of correlations was evaluated by fitting linear regression models, using Matlab’s fitlm function (Matlab 2019a).

### Models for mixed food patches

Consider mixtures of two strains, A and B, and let *N*_A_, *N*_B_ be the number of worms found experimentally in the two patches with pure bacterial cultures. For any other mixture, it computes the weighted arithmetic mean as *P*_*A*_*N*_A_ + (1 − *P*_*A*_)*N*_B_, and the weighted geometric mean as 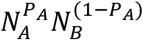, where *P*_*A*_ is the proportion of strain A in the mixture.

The model based on the sigmoidal rule assumes that the response to both strains follows the same sigmoid, with *H* = 146, *k* = 1.4 and with different attraction densities for each strain, *D*_attract,A_ and *D*_attract,B_. Therefore, if *D*_A_ and *D*_B_ are the optical densities of both pure strains (in all our experiments *D*_A_ = *D*_B_ = 0.5), their effective densities are *D*_A_*/D*_attract,A_ and *D*_B_*/D*_attract,B_. The effective density of a mixture with a proportion *P*_*A*_ of strain A and (1 − *P*_*A*_) of strain B will be *P*_*A*_ *D*_A_*/D*_attract,A_ + (1 − *P*_*A*_)*D*_B_*/D*_attract,B_. Using this effective density and **Equations 1 and 2**, we find the predicted proportion of worms in each food patch. Multiplying these proportions times the total number of worms provides the number of worms in each food patch.

## Acknowledgments

Some strains were provided by the CGC, which is funded by NIH Office of Research Infrastructure Programs (P40 OD010440). We are grateful to Céline Reyes for training and help with microscopy, Anna Mattout for training and help with DAPI staining, and members of IVEP team at the CRCA for comments on the manuscript.

## Author contributions

- Conceptualization – GM, JG, APE
- Data curation – AAA, GM, APE
- Formal analysis – AAA, GM, APE
- Funding acquisition – AAA, JG, APE
- Investigation – GM, AAA, LG, LML, AGE, MK, CR, APE
- Methodology – GM, CR, APE
- Project administration – JG, APE
- Resources – CR, APE
- Software – APE
- Supervision – GM, JG, APE
- Validation – AAA, APE
- Visualization – AAA, APE
- Writing – original draft – APE
- Writing – review & editing – AAA, GM, CR, JG, APE

## Supplementary Figures

**Figure S1:**
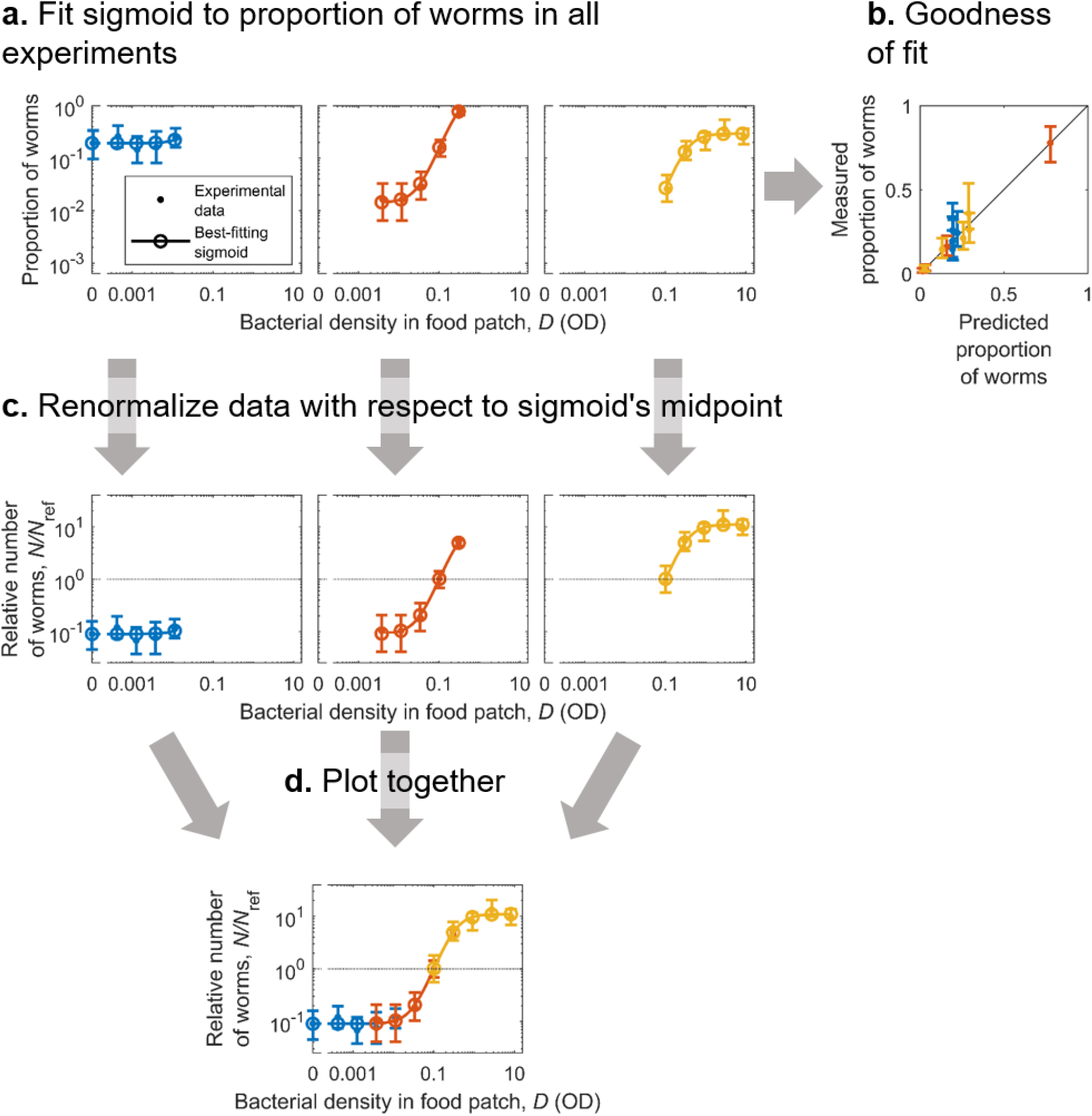
Illustration of data processing and normalization. Each complete sigmoid comes from several experiments, each of them covering part of the density range (in this case, 3 experiments with 5 food patches each; each of the three experiments is shown in one column and in a different color). **(a)** Proportion of worms present at each food patch in each experiment (dots with errorbars). These proportions add up to 1 for each experiment. Lines and circles: Best-fitting sigmoid. The three experiments (i.e. three columns) are described by the same sigmoid (**Equation 1** with a single set of parameters), but the sigmoid is normalized for each experiment using **Equation 2. (b)** Measured proportions versus predicted proportions, for all food patches of the three experiments. The black line corresponds to a perfect prediction. **(c)** Normalized number of worms, *N/N*_ref_, where *N* is the number of worms in each food patch, and *N*_ref_ is the number of worms in a virtual reference patch (**Equation 4**). **(D)** Same as (C), but with all data in the same plot.

**Figure S2:**
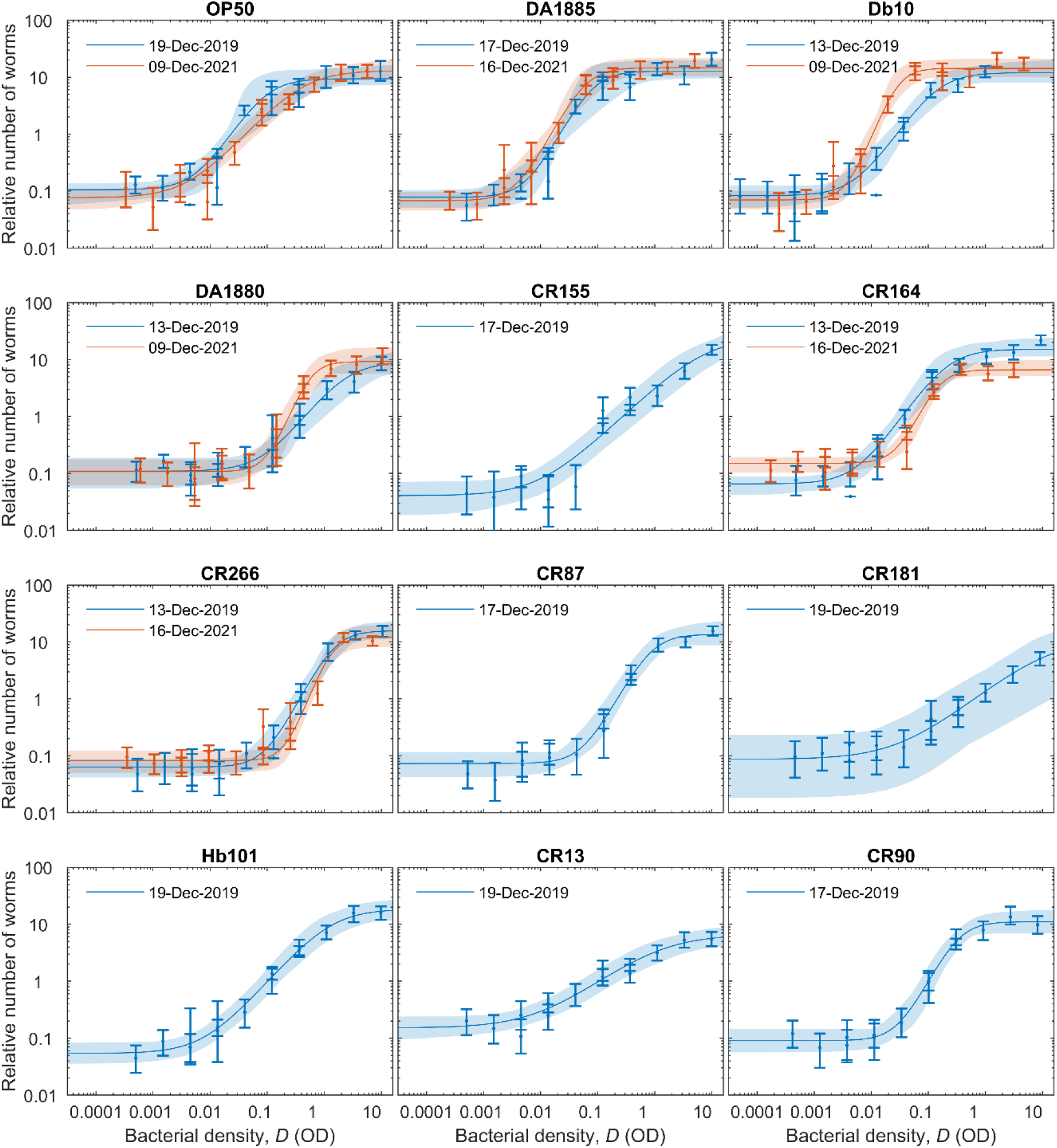
Sigmoids for all strains. Relative number of worms found at each food patch, as a function of bacterial density in the food patch (*D*, measured in Optical Density, OD). Dots: Experimental data; errorbars show the 95% confidence interval. Lines: Best-fitting sigmoid. Semitransparent patches: 95% confidence interval for the fitted sigmoids. Six strains were measured twice, and the lockdown due to the COVID-19 pandemic imposed a two-year gap between them, a period during which the laboratory moved to a new building. As a result, both replicates were performed in different conditions: 2019 experiments were performed in an incubator and the main experimenter was LG, while 2021 experiments were performed in an environmental room and the main experimenter was AAA. All other experimental parameters were kept as equal as possible. We chose to show these results rather than repeating the experiments in the same conditions to highlight the robustness of our results: While two strains (Db10 and CR164) show significant differences across the two replicates, we found remarkable reproducibility given the experimental differences. Results shown in **Figures 1** and **3** of the main text correspond to the 2019 experiments.

**Figure S3:**
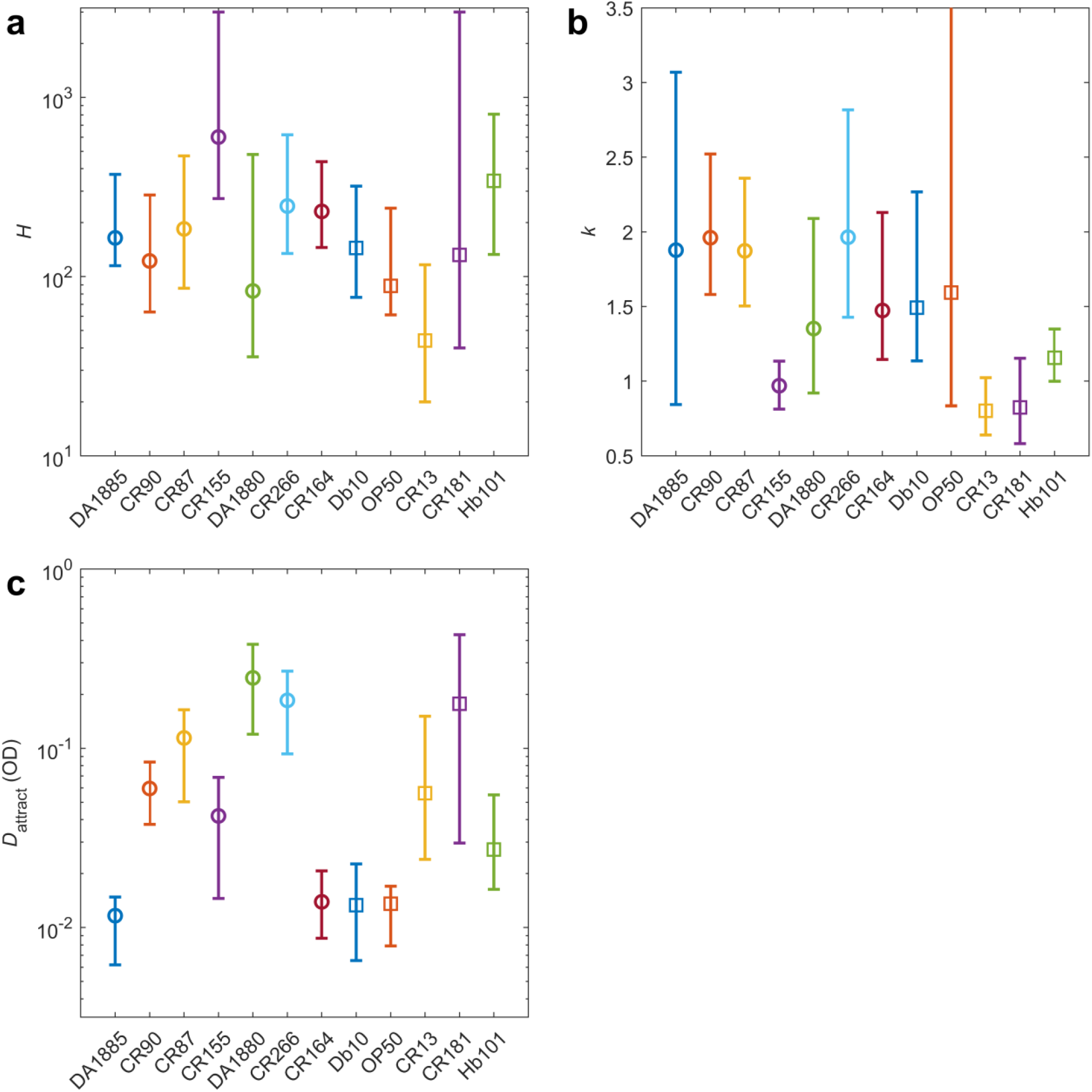
Sigmoid parameters for all strains. **(A)** Best-fitting value of *H* for each bacterial strain. **(B)** Best-fitting value of *k* D0 for each bacterial strain. **(C)** Best-fitting value of the attraction density (*D*_attract_) for each bacterial strain. All errorbars show the 95% confidence interval, calculated via bootstrapping.

**Figure S4:**
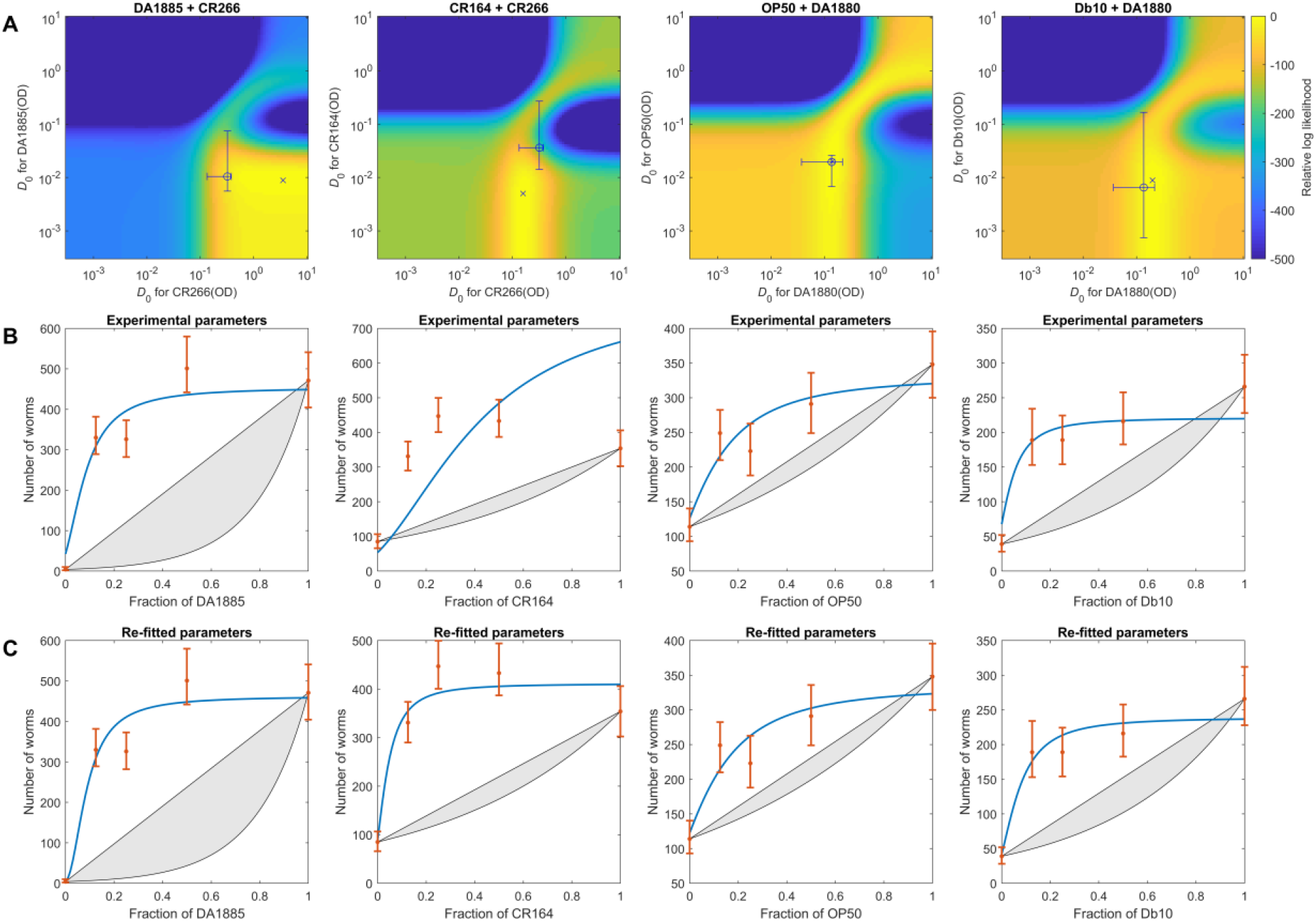
Results from mixed drops. Each column corresponds to one pair of strains. **(A)** Goodness of fit of our prediction to the experimental data, for each possible value of *D*_attract_ for each strain (hotter color means better fit). Circle with errorbars: Experimental *D*_attract_, obtained from the sigmoidal fit to an independent experiment (orange sigmoids in **Figure S2**). Cross: Values of *D*_attract_ that best reproduce the mixed-drop experiments. In the last two pairs, the experimental values are very close to the best-fitting one. In the first pair the optimal value is far from the experimental one, but the experimental values are still on the region of excellent fit (yellow region). In the second pair the fit is not good: The experimental values fall outside the good fitting region. This pair involves strain CR164, which also gave a sigmoid significantly different to the one originally measured (see **Figure S2**). Therefore, it’s likely that experimental variability was responsible for this mismatch, but we cannot exclude other factors. **(B)** Number of worms at each food patch, as a fraction of one of the strains. Dots: Experimental results (errorbars show the 95% confidence interval). Red line: Prediction from our model, using the experimentally measured *D*_attract_ (circle in box A). Gray patch: Prediction of the null model. **(C)** Same as B, but prediction uses the best-fitting values of *D*_attract_ (cross in box A).

**Figure S5:**
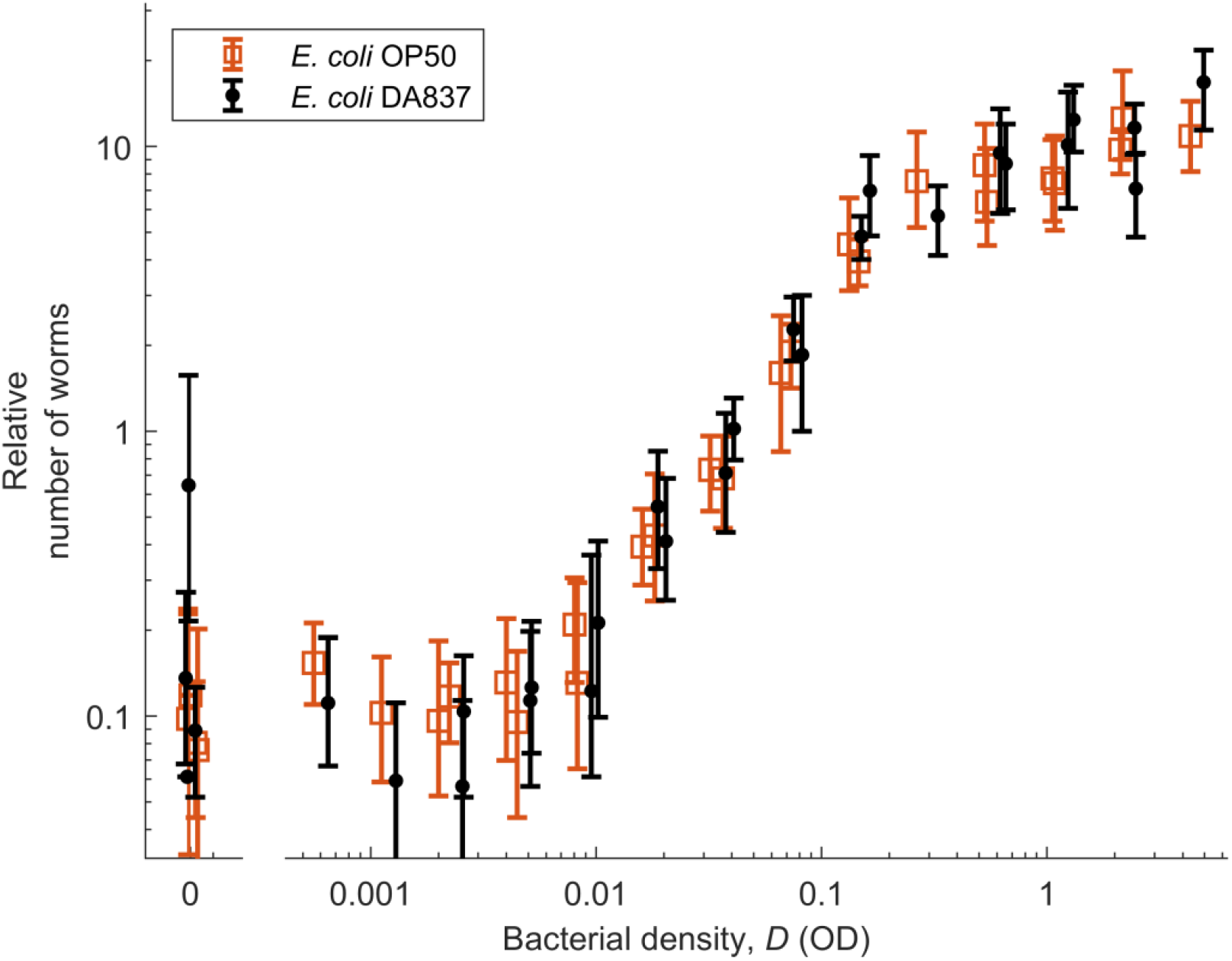
*C. elegans* shows the same response to OP50 and DA837. Relative number of worms found at each food patch, as a function of bacterial density in the food patch (*D*). Squares: *E. coli* OP50. Dots: *E. coli* DA837. Errorbars show the 95% confidence interval, computed via bootstrapping.

**Figure S6:**
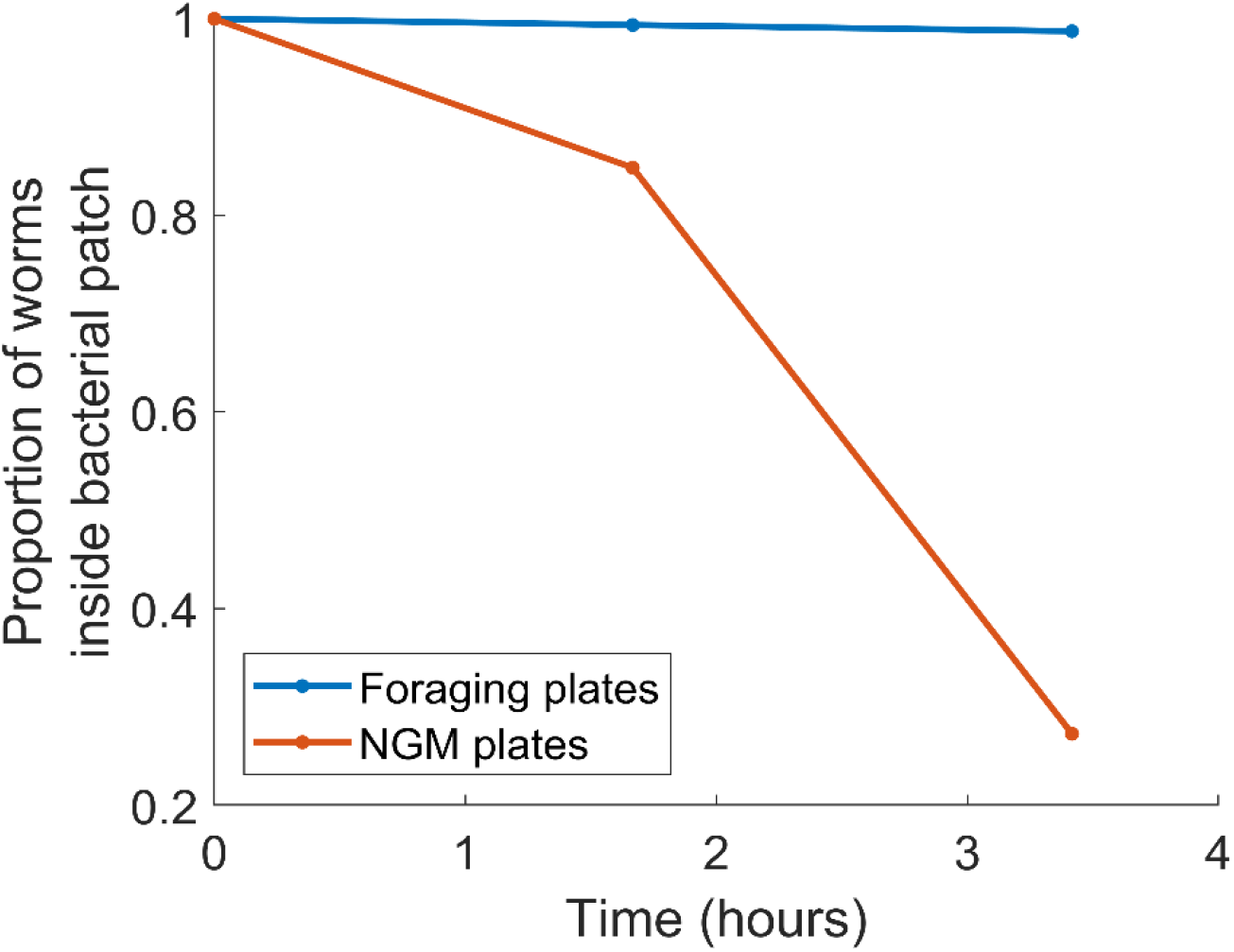
Avoidance of *Serratia marcescens* requires active bacterial growth. We placed a 40 microliter drop of saturated overnight culture of *S. marcescens* (Db10) at the middle of either an NGM plate (where bacteria can grow) or a foraging plate (where bacteria cannot grow due to lack of nutrients and presence of bacteriostatic antibiotics). The next day, we placed around 30 young adult worms (48-hour old) at the center of the food patch. The figure shows the proportion of worms that remained inside the food patch as a function of time, for both treatments. These results are consistent with previous studies, which found strong avoidance of *S. marcescens* on NGM plates.^36^

